## Supplementary figures and images for "ScalaFlux: a scalable approach to quantify fluxes in metabolic subnetworks"

### S1 Fig

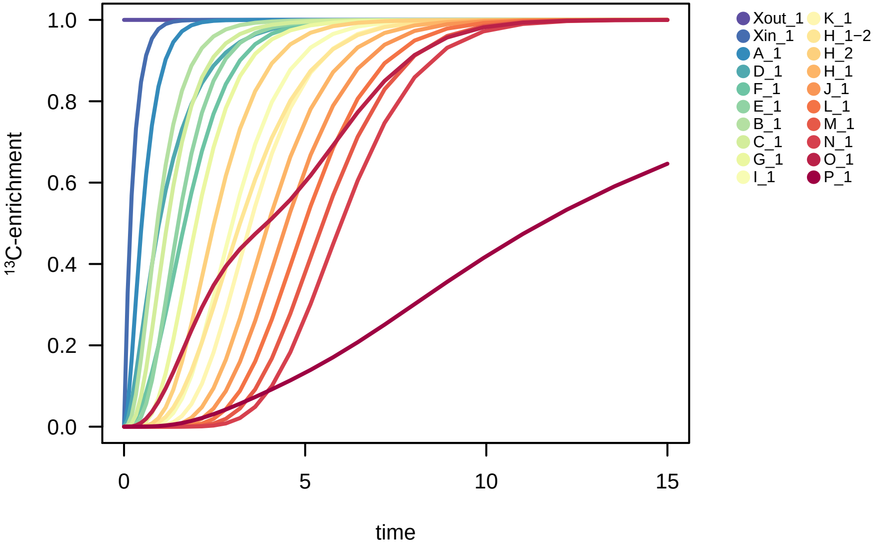

### S2 Fig

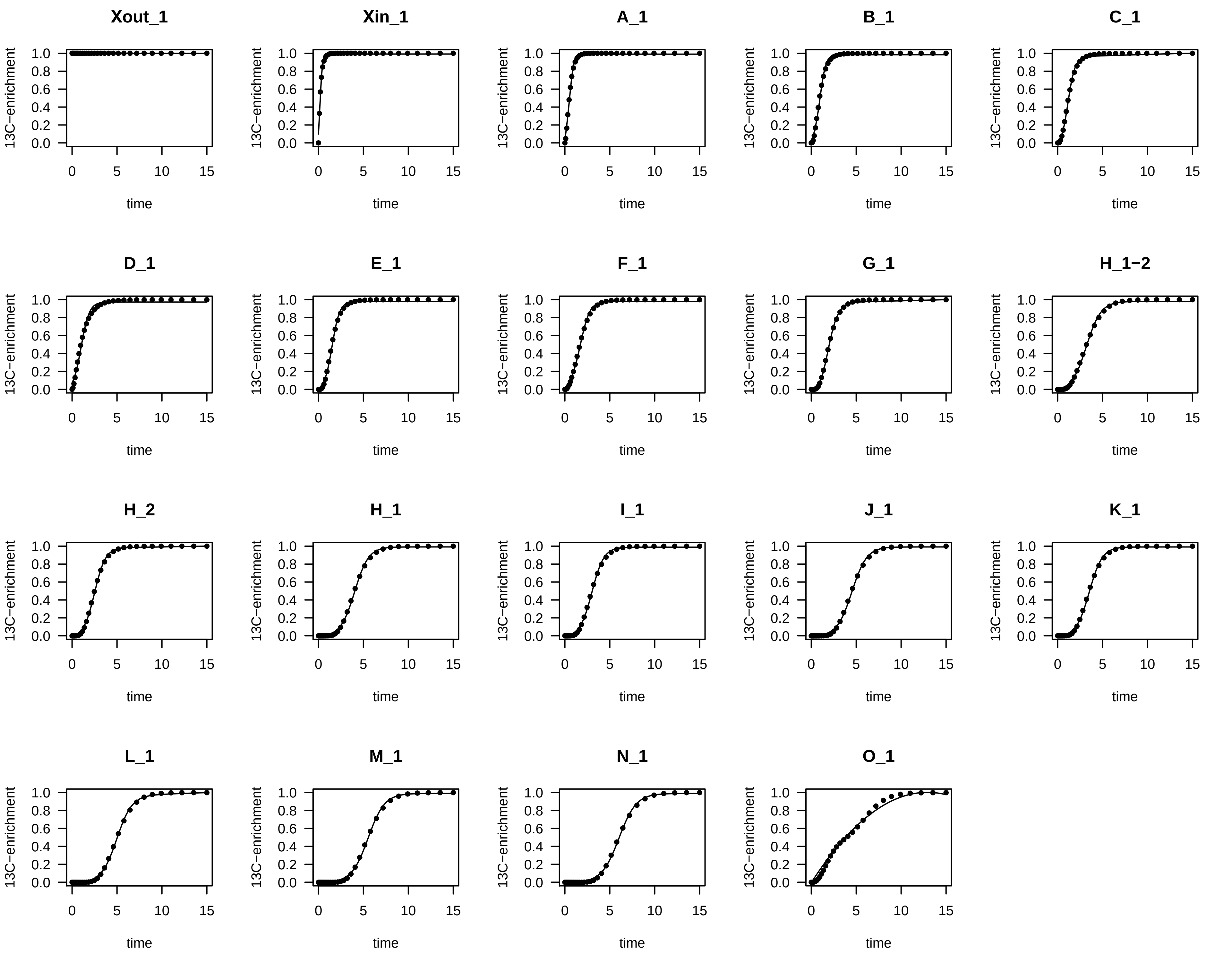

### S3 Fig

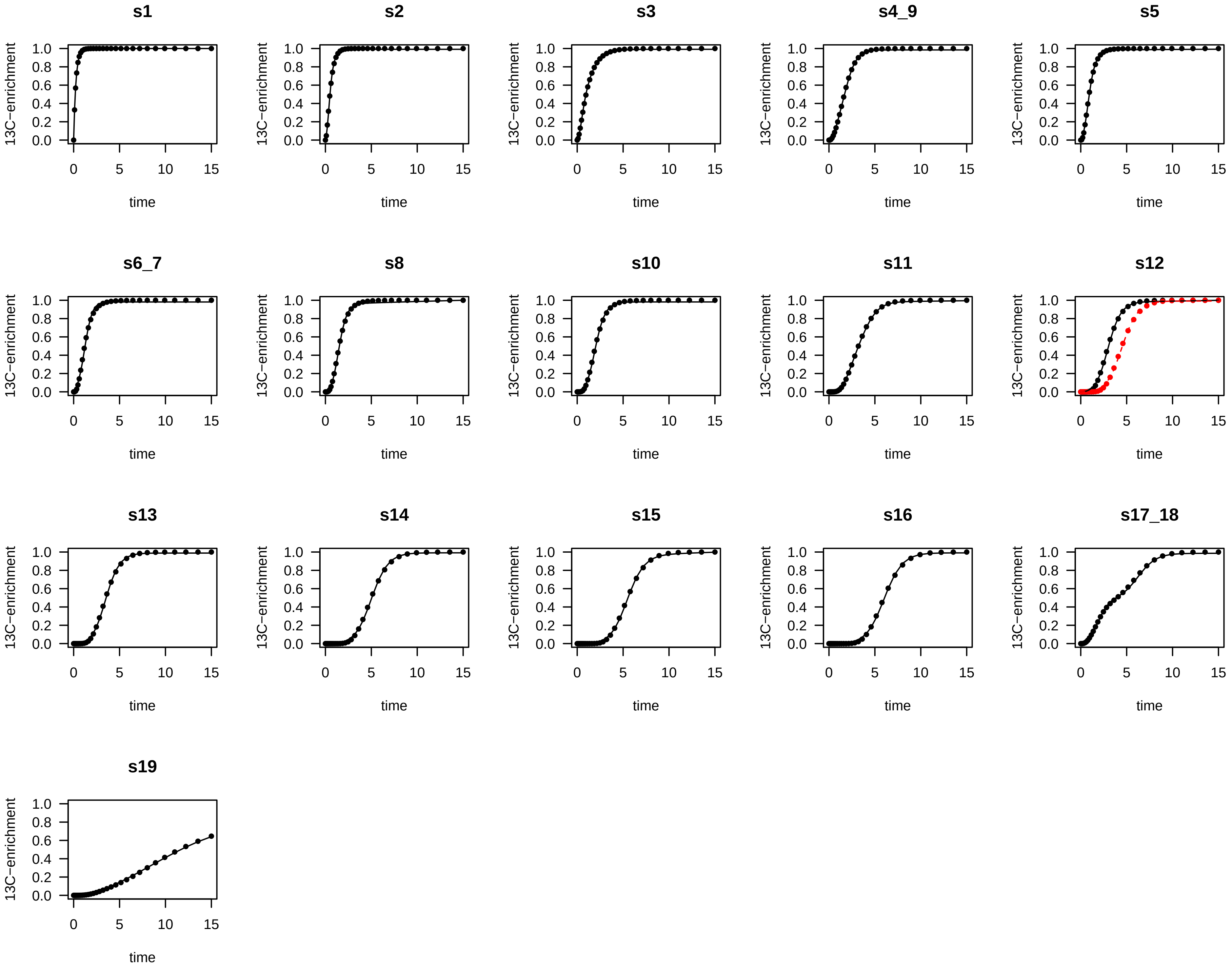
